# Talking Brains: Neonatal Aperiodic Activity in Resting State EEG Relates to Later Communicative Abilities

**DOI:** 10.64898/2026.09.05.749585

**Authors:** Michaela Reimann-Ayiköz, Mohamed S. Ameen, Eva Reisenberger, Cristina Florea, Jasmin Preiß, Monika Angerer, Manuel Schabus, Dietmar Roehm, Dominik Heib, Claudia Männel

## Abstract

Communicative abilities acquired over the first year of life are foundational for language development. Understanding the neural mechanisms underlying these abilities can explain individual differences and improve predictions of later language outcomes. While prior research on neural markers of language development has focused predominantly on oscillatory neural activity, the underlying aperiodic component of the neural signal may provide further insights into cortical maturation by indirectly reflecting variation in neural excitation/inhibition (E/I) balance. In neurodevelopmental conditions, both reduced and elevated aperiodic activity are linked to atypical development, and initial evidence from at-risk cohorts also relates aperiodic activity to language outcome. Moving beyond clinical populations, the present study examined whether aperiodic activity during neonatal resting-state EEG relates to 12-month communicative abilities in typically developing infants. Aperiodic offset and exponent were extracted from resting-state EEG recorded two weeks after birth in *n* = 50 neonates using the FOOOF algorithm. At 12 months, communicative abilities were assessed with the parent-report CSBS-DP Infant-Toddler Checklist. Quadratic regression analyses showed an inverted U-shaped relationship between the neonatal aperiodic exponent and communicative abilities: both lower and higher exponent values were associated with poorer outcomes, with intermediate values linked to better abilities. No robust association was found for the offset. These findings suggest that aperiodic activity captures variability in early communicative development, with an optimum at intermediate levels, consistent with a balanced E/I state. The scope of aperiodic brain activity as a developmental marker may thus extend beyond clinical populations to predicting communicative development in healthy infants.

## Introduction

In the neonatal brain, most neurons have already been generated, major white-matter tracts are in place, and white-matter structural and sensorimotor functional networks are largely established (for reviews, see [1,2]). Nevertheless, the first year of life marks a remarkable period in human brain development, characterized by rapid learning and the fastest structural and functional changes across the lifespan (for reviews, see [2,3]), with substantial growth in overall brain volume, progressive myelination, expansion of cortical thickness and surface area, and the emerging organization of functional networks [4,5] (for reviews, see [1,6]).

As the brain matures, neural oscillatory activity changes markedly across the first year of life, as reflected in a substantial shift in the neural power spectrum. At birth, the EEG is characterized by slow-frequency activity, with delta (1-3 Hz) and theta (3-6 Hz) frequencies dominating the spectrum [7] (for reviews, see [8,9]). With increasing age, neural power extends towards higher frequencies, marked by the emergence of an infant alpha rhythm (6-9 Hz) in the first half of the first year and subsequent increases in faster frequencies [10–12].

Understanding oscillatory activity provides insights into how the brain functions [13,14] (for a review, see [15]). In infancy, variations in oscillatory power have been linked to the development of language and communicative abilities (e.g., [16–20]). Yet findings vary with respect to the frequency bands examined (the boundaries of which are not uniformly defined in infancy; for a review, see [9]), the developmental stage assessed, and the direction of the effect. While most studies report that higher oscillatory power is associated with higher language abilities [16,17,19,20], Piazza and colleagues [18] observed the opposite pattern, with increased power in children at familial risk for language learning impairments compared with controls. These inconsistencies point toward a more nuanced evaluation of the neural signal, as focusing solely on band-specific oscillatory measures may provide limited differentiation of the underlying spectral contributions that may play a role in the development of language abilities.

Conventional oscillatory power analyses capture periodic activity alongside contributions from the aperiodic component, yet aperiodic activity has often been treated as background noise. More recent work, however, has demonstrated that the aperiodic component itself reflects physiological properties of cortical activity that may be relevant for functional brain development (see also [21,22]). The aperiodic signal can be parametrized into two components, that is, offset and exponent. The offset reflects a broadband shift in power across frequencies and has been functionally linked to neuronal population spiking [23] and hemodynamic activity, potentially reflecting arousal-related cortical activation [24]. The exponent indicates the slope of the 1/f-like spectrum and thus describes how steeply neural power decreases with increasing frequency [21,25]. A higher exponent, that is, a steeper slope of the aperiodic component, indicates relatively greater power at lower frequencies compared with higher frequencies, whereas a lower exponent indicates a flatter spectral profile. The exponent thus characterizes the temporal organization of ongoing neural activity [26], which may partly reflect how excitatory and inhibitory synaptic currents contribute to field potentials over time (for a review, see [27]). Building on this relationship, the exponent has been proposed as an indirect measure of excitation/inhibition (E/I) balance [28]. E/I balance describes the dynamic coordination between excitatory and inhibitory synaptic activity that generates local network activity and shapes the strength, timing, and gain of cortical responses [29] (for a review, see [30]). Thus, the aperiodic exponent may provide further insights into how cortical networks are organized across development (see also [22]).

Aperiodic activity has been reported to change systematically with age (e.g., [31,32]). In the prenatal period, the aperiodic exponent increases with advancing gestational maturity, as indicated by positive associations with both post-conceptual age [33] and gestation duration [34]. However, no such association was found for the aperiodic offset [34]. Across infancy, findings on the developmental trajectory of aperiodic activity remain inconsistent. Some studies described the aperiodic exponent decreasing throughout the first year of life [32,35], with no age-related changes in the offset [35]. In contrast, Wilkinson and colleagues [36] found increases in both aperiodic offset and exponent during infancy. During childhood and adolescence, both offset and exponent decrease with age [31,37], and evidence also points to non-linear developmental trajectories [38]. Thus, current reports vary on the direction of developmental changes in aperiodic activity but together indicate that aperiodic activity captures differences in the maturation of cortical organization, while the functional implications of these processes remain to be fully understood.

Despite its potential relevance in early neurodevelopment, aperiodic activity has so far received little attention in research on language development. Only a few studies have reported altered aperiodic activity in the pathophysiology of neurodevelopmental disorders. Compared to healthy controls, steeper slopes have been reported in children with autism spectrum disorder (ASD) [39–41] and Rett syndrome [41], flatter slopes in children with Down syndrome [42], and mixed findings, both steeper and flatter slopes, for children with attention-deficit/hyperactivity disorder (ADHD) [43–45]. Notably, many of these neurodevelopmental conditions are characterized by communication and language difficulties, suggesting that aperiodic activity may relate to emerging language abilities. Here, exploratory longitudinal analyses by Wilkinson and colleagues [46] provided initial evidence that larger increases in aperiodic offset and exponent in the first year of life are associated with lower language abilities at 18 months in children with and without a family history of ASD. In contrast, no significant associations between aperiodic offset or exponent and language development were found in children with Down syndrome at age one to four years [42]. In older children aged six to 17 years, both with and without a diagnosis of ASD, age-related patterns of E/I balance have been related to language ability [47]. Specifically, an inverse U-shaped trajectory of E/I balance across groups was associated with lower language abilities and a U-shaped trajectory with higher language abilities. Together, these studies provide initial insights into the role of aperiodic activity in language development, but have predominantly focused on clinical or at-risk populations. Against this background, examining aperiodic parameters in a typically developing population could extend previous research into oscillatory markers of language development and advance our understanding of early physiological mechanisms underlying development.

In the current study, we examined the relationship between neonatal aperiodic activity and communicative abilities at 12 months of age as important precursors of later language development (see also [48]). Specifically, we focused on the aperiodic exponent and offset derived from resting-state EEG recordings obtained two weeks after birth, given that resting-state EEG is particularly informative for assessing intrinsic brain activity in the absence of task-related demands. We hypothesized a non-linear quadratic relationship between neonatal aperiodic activity and later communicative abilities, based on previous findings in neurodevelopmental disorders, suggesting that deviations in either direction may be associated with neurodevelopmental conditions (e.g., [39,41,42]). Given the limited evidence in typically developing infants and children, this hypothesis is further informed by theoretical work on neural communication [49] across development, aging, and psychiatric and neurological disorders, proposing that excessively steep or flat aperiodic 1/f slopes may reflect altered coordination of neural activity. Correspondingly, for our non-clinical sample, we expected the aperiodic exponent to be associated with less favorable communicative outcomes at both relatively high and relatively low levels. Although the role of the aperiodic offset has received less attention so far, we hypothesized a similar pattern as for the exponent, as both aperiodic signals are closely intertwined (see also [22]).

## Methods

### Participants

Data reported in this study were collected as part of a longitudinal project, funded by the Austrian Science Fund (FWF), following *n* = 67 mother-infant dyads from the 34th week of gestation until 12 months after birth. The aim of the project was to examine the early foundations of neurodevelopment and to evaluate how prenatal and postnatal factors contribute to early development. The Ethics Committee of the University of Salzburg approved the study procedures (EK-GZ: 12/2023, positive vote on 23rd of January, 2014, Addendum: 17th November, 2021). Prior to the study, all mothers received detailed information about the study procedure and provided written consent. Inclusion criteria required pregnant women to be at least 18 years old, to speak German as their first language, and to experience no pregnancy complications, except for transient findings in *n* = 3 women (i.e., resolved placenta previa; cervical insufficiency without further clinical relevance; hyperemesis and abnormal nuchal translucency with unremarkable follow-up NIPD result). Additionally, smoking and alcohol consumption during pregnancy were exclusion criteria, with one dyad being excluded. Infant inclusion criteria were as follows: no developmental delay in cognitive or language domains at 12 months according to pediatric evaluation, a gestational age of at least 37 weeks (with one dyad being excluded due to preterm birth), German as the primary environmental language, no hearing impairment, and no history of language impairments or autism in first- and second-degree relatives. Demographic maternal and child-specific information was collected via a study-specific questionnaire (see Table 1 for the final sample of *n* = 50).

**Table 1.** Demographic and child-specific characteristics of the final sample (*n =* 50)

| Variable |  |
|---|---|
| **Maternal and family characteristics** |  |
| Maternal age at birth (in years), mean (SD) [range] | 32.40 (4.59) [20-40] |
| Maternal relationship status at the 34th GW (%) |  |
| Married to the child’s father | 52 |
| In a relationship with the child’s father | 44 |
| Single | 4 |
| Professional qualification of mothers (%) |  |
| Without vocational qualification | 2 |
| Completed vocational training | 20 |
| Higher vocational education | 14 |
| University degree (or higher) | 64 |
| **Infant characteristics** |  |
| Sex (% female) | 52 |
| Gestational age at birth in weeks, mean (SD in days) [range] | 40+1 (10) [37-42] |
| Delivery mode (%) |  |
| Vaginal delivery | 64 |
| Operative vaginal delivery (forceps or vacuum) | 10 |
| C-section | 26 |
| Variable |  |
| Birth weight (in g), mean (SD) [range] | 3350 (398) [2540-4740] |
| Birth height (in cm), mean (SD) [range] | 51.52 (2.38) [48-58] |
| Apgar-score (max. 10),10 minutes after birth, mean (SD) [range] | 9.88 (0.59) [6-10] |
| Age at two-weeks EEG assessment, mean (SD) [range] | 16.40 d (3.93) [12-33] |
| Head circumference (in cm) at EEG assessment, mean (SD) [range] | 36.11 cm (1.20) [34-39] |
| Age at 12-months communicative skills assessment, mean (SD) [range] | 12.33 M (0.47) [11.43-13.93] |

Figure 1 provides an overview of the participant inclusion procedure, with the final sample comprising 50 datasets (26 females). We initially started with a sample of *n* = 65 mother-infant dyads to obtain both neonatal EEG data at two weeks after birth and maternal reports of communication abilities at 12 months postpartum. For *n* = 4 children, maternal reports were not available, and following EEG preprocessing, data from *n* = 11 children had to be excluded (see Figure 1 for criteria). Please note that all subsequent analyses refer to the final sample (*n* = 50), unless stated otherwise.

**Figure 1.**
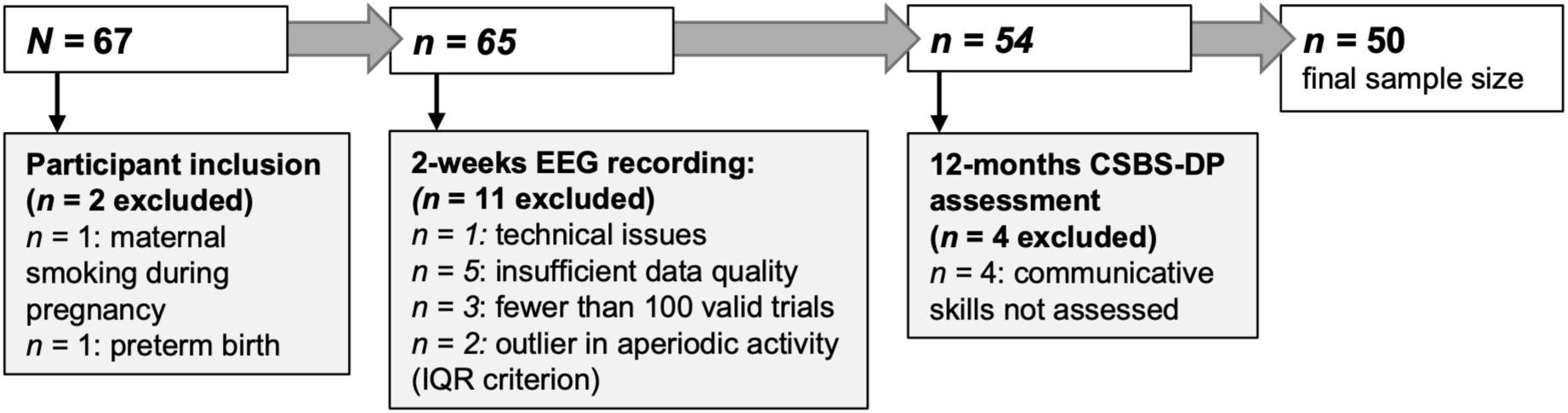
Flowchart of sample inclusion and exclusion criteria across study assessments, resulting in a final sample of *n* = 50 infants. *Note.* IQR = Interquartile range; CSBS-DP = Communication and Symbolic Behavior Scales Developmental Profile Infant-Toddler Checklist

### EEG acquisition and procedure

EEG data were collected in neonates approximately two weeks after birth (*M*_age_ = 16.40 d, *SD* = 3.93). The experimental paradigm comprised seven silent baselines (see Figure 2), providing 12 minutes and 45 seconds of resting-state EEG data. These baselines were embedded in a larger protocol, with a total recording duration of 29.5 minutes (for a detailed description, see [50]): the experiment was initiated with a 30-second silent baseline (not available in the first two participants, as the paradigm was slightly modified thereafter), followed by a 135-second beep-tone sequence. Thereafter, five conditions of nursery rhymes were presented, alternating with five 120-second silent baselines, followed by a final 135-second silent baseline.

**Figure 2.**
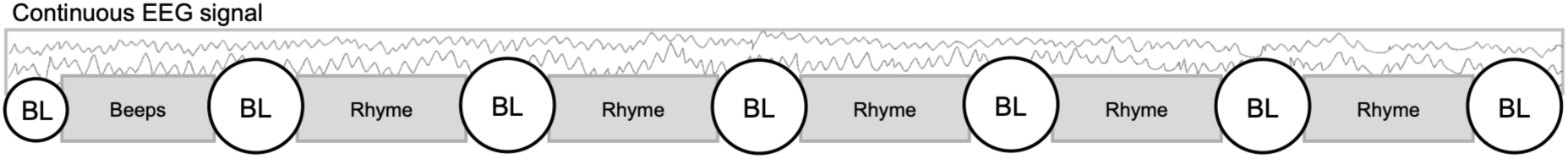
Overview of the EEG paradigm. *Note.* The EEG paradigm comprised an initial 30-s silent baseline (BL), followed by a 135-s beep-tone sequence, five nursery-rhyme conditions alternating with five 120-s silent baselines, and a final 135-s silent baseline. Analyses were based exclusively on the silent baseline periods (12 min 45 s).

The data were collected either at the families’ homes (*n =* 41) or at the infant laboratory at the university (*n =* 9), depending on parental preference. To maximize comfort, the infant’s position was kept flexible during data collection, with most infants held in their caregiver’s arms. Although most infants were asleep during testing, some variability in sleep-wake state was observed within and across participants. To account for this variability, we conducted supplementary analyses (see supplementary information (SI) S5) using only periods during which infants were asleep (*n =* 45).

### EEG data acquisition

Continuous EEG was recorded at a sampling rate of 1000 Hz using a high-density 124/128-channel HydroCel Geodesic Sensor Net (Electrical Geodesics, Inc. [EGI], Eugene, OR, USA) connected to a GES 400 amplifier (EGI, Eugene, OR, USA). Data were acquired using the NetStation software (version 5.4.2, release r29917). Before recording, electrode impedances were checked and kept below 50 kΩ. The EEG was referenced online to the vertex electrode (Cz).

### EEG data analysis

The data were preprocessed using the EEGLAB toolbox 2024.2 [51] and a custom MATLAB code (R2023b, Update 10, 23.2.0.2859533). The preprocessing procedure was based on Reisenberger and colleagues [52], with several modifications to accommodate the specific requirements of the current analysis.

### EEG pre-processing and rejection criteria

Following a visual inspection of the data, five datasets were excluded from the EEG analysis due to insufficient data quality, resulting from movement- and handling-related artifacts. One additional dataset was excluded due to file damage. The first preprocessing step involved removing the outermost electrode rows, leaving 91 electrodes available for further analysis (see SI, Figure S1 for EEG layout). Next, data were band-pass filtered from 1 to 45 Hz using a Hamming-windowed non-causal sinc Finite Impulse Response filter implemented in *pop_eegfiltnew* (filter order: 3,301; −6dB cut-off frequencies: 0.5 and 45.5 Hz). Segments containing pronounced artifacts, such as abrupt data offsets, were manually removed from the continuous data to ensure sufficient data quality for further data processing. As a next step, flat channels were identified using *clean_artifacts* with a FlatlineCriterion of 5 seconds, and noisy channels were identified using a fixed peak-to-peak amplitude (> 80,000), as well as *pop_rejchan* based on kurtosis (threshold = 5), probability (threshold = 5), and spectral criteria (threshold 25 within 2-40 Hz). The PREP pipeline [53] was then applied to clean the data further (badTimeThreshold = 0.01; highFrequencyNoiseThreshold = 5; robustDeviationThreshold = 5; ransacOff = true; interpolateChannels = false) and to compute and apply a robust average reference while excluding the channels that had been identified as flat or noisy. These channels were then interpolated using spherical interpolation with *pop_interp*, with a predefined threshold of 25% for the proportion of interpolated channels within the region of interest. Note that none of the participants reached this threshold (*M*_interpolated channels_ = 0.82/24 channels, *SD* = 1.00, ranging from 0 to 4 channels). Finally, an extended ICA was performed for artifact correction via *runica*. Independent components were automatically classified using *ICLabel* [54], and artifact-related ICA components (*M*_number of rejected components_ = 5.54, *SD* = 2.99, ranging from 0 to 12 components out of *M* = 78.54 components, *SD* = 7.40, range = 50-89) were rejected based on established probability thresholds (≥ .80 classification probability).

### Extraction of aperiodic components from resting-state EEG

All analyses of aperiodic spectral activity were performed in FieldTrip [55]. First, continuous resting-state EEG data were segmented into six-second epochs with a one-second recurring window (*M*_epochs_ = 385.48, *SD* = 149.30, range = 113-627). To minimize edge artifacts potentially introduced by filtering (see also [56]), the first four seconds of the initial baseline and the last four seconds of the final baseline were excluded. Additionally, the first six seconds following each stimulation were omitted to minimize post-stimulation effects. Epochs identified as artifactual were automatically rejected using *perform_epoch_rej* (default threshold = 1000, maxrej = 90). Only data from participants with at least 100 artifact-free resting-state epochs were retained for further analysis (*M*_clean epochs_ = 358.98, *SD* = 141.60, range = 102-627), resulting in the exclusion of *n =* 3 additional participants.

For the estimation of aperiodic spectral activity, power spectra were computed for all retained epochs using the multitaper method (cfg.method = ‘mtmfft’; cfg.pad = ‘nextpow2’). To separate aperiodic and periodic components of the spectrum, spectral parameterization was performed with the FOOOF algorithm (i.e., Fitting Oscillations and One-Over-F, https://fooof-tools.github.io/fooof), a MATLAB-based tool in FieldTrip [21]. This algorithm models the power spectrum as an aperiodic component and Gaussian peaks representing periodic activity. Previous infant EEG studies have applied FOOOF to a variety of frequency windows, from narrower frequencies such as 1-10 Hz [32,40] and 1-20 Hz [35] to broader ranges extending up to 55 Hz [36,39]. In our data, estimates of the aperiodic exponent and offset derived from ranges 1-15 Hz and 1-30 Hz were highly correlated (see SI, Figure S2). We therefore selected the broader 1-30 Hz range for all analyses, in line with recommendations that wider fitting ranges improve the robustness and interpretability of aperiodic parameter estimates by better capturing the overall spectral shape, while minimizing edge-related biases [57]. For spectral fitting, the following parameter settings were used: peak_width_limits = [0.5, 12.0], max_n_peaks = 5, peak_threshold = 2.0, min_peak_height = 0, aperiodic_mode = ‘fixed’, and verbose = true. For each participant and electrode, the aperiodic exponent and offset values were extracted from the fitted aperiodic signal (see Figure 3).

**Figure 3.**
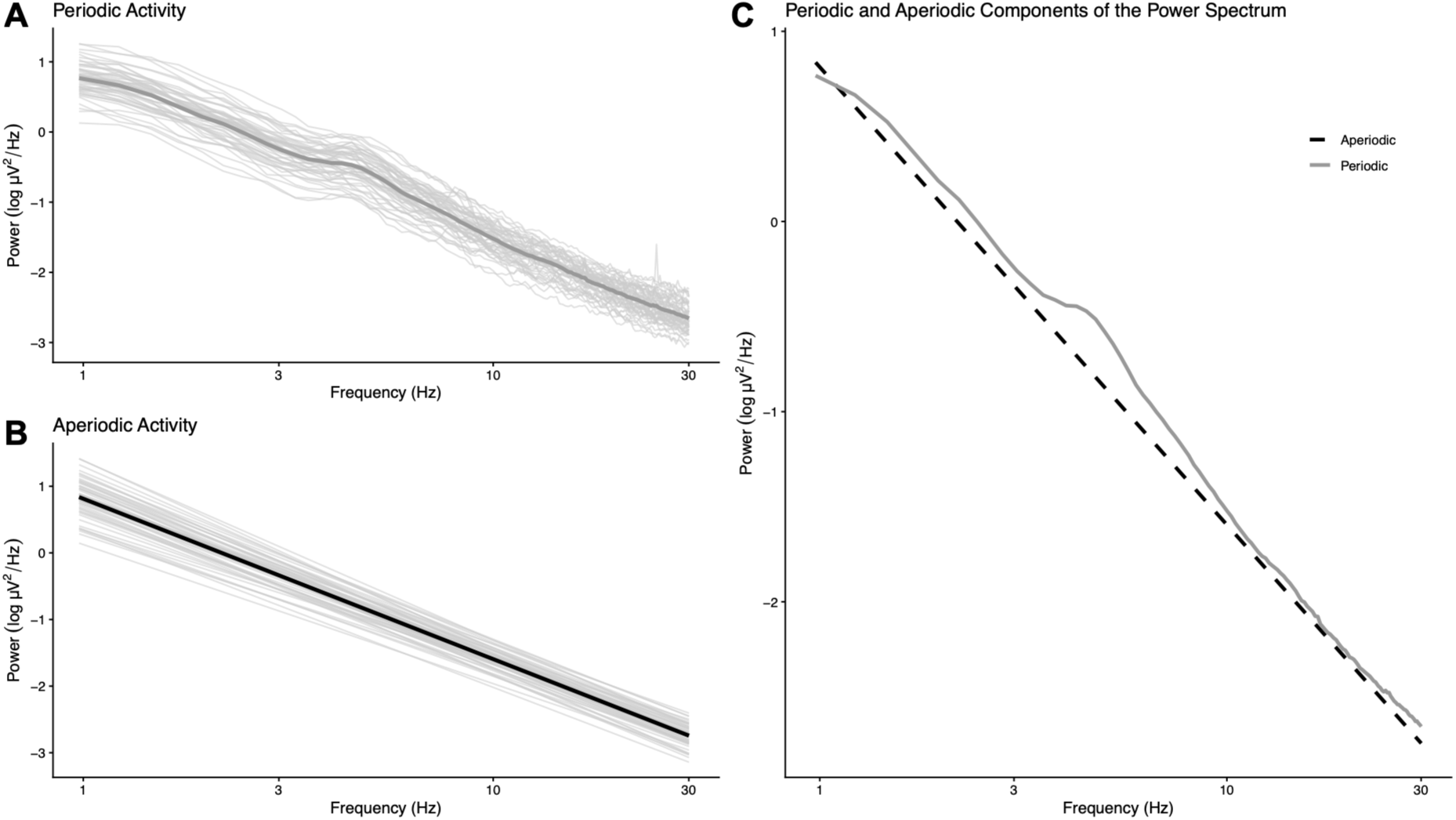
Illustration of the decomposition of the power spectrum across the 1 to 30 Hz range for the final sample (*n* = 50), showing (A) periodic activity, (B) aperiodic activity, and (C) the combined power spectrum.

Based on a defined quality criterion of FOOOF, only electrode-wise model fits with R^2^ > 0.9 were included (*M*_R_squared_ = 0.994, SD = 0.006, ranging from 0.923 to 0.999) in the analysis. In one participant, one electrode fell below this threshold and was excluded, but the dataset was retained because >95% of the data remained. For the statistical analysis, aperiodic parameters were averaged across 24 frontal electrodes for each participant (see SI, Figure S1 for EEG layout), providing a more stable and robust estimate by reducing variability across individual electrodes. The frontal region was selected based on previous findings on the relation of neural activity (periodic and aperiodic) and language development (e.g., [16,17,46]).

### Communicative abilities

At 12 months of age, children’s communicative behavior was assessed using the German version [58] of the parental questionnaire *Communication and Symbolic Behavior Scales Developmental Profile (CSBS-DP) Infant-Toddler Checklist*, with norms based on 2,000 children aged six to 24 months [48]. The questionnaire measures early communicative development and helps identify children at risk for communication and language difficulties. The *communication composite* covers emotion and eye gaze as well as the use of communication and gestures; the *expressive speech composite* includes the production of speech sounds and words; and the *symbolic composite* comprises language comprehension and object use. In total, the questionnaire consists of 24 items, with each item scored from 0 to 2, 0 to 3, or 0 to 4 points, depending on the number of response options. In the present study, we analyzed the total score (0-57), calculated as the sum of the three composite scores. Thereby, higher scores reflect more advanced communicative abilities, with scores below 28 in children aged 12 months being considered indicative of risk for communication difficulties (*n =* 3, all male).

### Statistical analyses

All statistical analyses were conducted in RStudio 026.01.0+392 [59]. Multiple linear regression models were used for statistical analyses, with 12-month CSBS-DP Infant-Toddler Checklist total score as the outcome variable and the aperiodic components as explanatory variables. We ran separate models for offset and exponent to avoid multicollinearity, as they were highly correlated (*r_spearman_ =* .80, *p <* .001). Both linear and quadratic terms of the aperiodic component were entered into the models, allowing us to test the hypothesized non-linear relationship while controlling for the linear effect. Before statistical analysis, two datasets were excluded as outliers based on the interquartile range (IQR) criterion for the aperiodic offset and exponent. To reduce multicollinearity between the linear and quadratic terms, aperiodic components were z-standardized prior to inclusion in the regression models. Given that age at 12 months (*r_spearman_ =* .08, *p =* .57) and gestational age at birth (*r_spearman_ =* xys.14, *p =* .33) were not significantly correlated with the communication outcome variable, age measures were not included as covariates in the model. Although there were no significant sex differences in communicative abilities in the full sample (t(40.81) = 1.37, *p =* .18), analyses were controlled for sex, as all three children at risk of communication difficulties were male.

Overall, regression assumptions were met, including the normality of the residuals, homoscedasticity, and the absence of multicollinearity (VIF <1.2). Given the presence of influential cases, the models were re-estimated using robust regression [60] to assess the robustness of the findings.

We conducted control analyses with gestational age as predictor to examine whether our findings were attributable to gestational age rather than aperiodic activity (see SI, Table S4), given that the aperiodic exponent has previously been linked to gestational age [34] and consistent with our data (exponent: *r_spearman_ =* .36, *p =* .01; offset: *r_spearman_ =* .21, *p =* .15). Moreover, to account for potential effects of sleep-wake state, we conducted additional analyses including only EEG periods during which the infants were asleep (for details, see SI Tables S5a and S5b). As a further comparison, associations between absolute delta and theta power and communicative abilities were examined using conventional band-power analysis (for details, see SI Tables S6a and S6b). The significance level was set at *p* = .05.

## Results

Table 2 provides descriptive statistics for the aperiodic offset and exponent assessed at two weeks postpartum and for communicative skills at 12 months, examined with the CSBS-DP Infant-Toddler Checklist for the total sample and divided by sex. No statistically significant sex differences were observed in any of these variables.

**Table 2.** Descriptive Statistics for neonatal aperiodic offset and exponent, and communicative abilities at 12 months for the total sample (*n* = 50) and separately by sex.

|  | Final sample<br>(*n* = 50) | Males<br>(*n* = 24) | Females<br>(*n* = 26) | Sex comparison |
|---|---|---|---|---|
| Aperiodic offset | 0.81 (0.29)<br>[0.12-1.39] | 0.85 (0.35)<br>[0.12-1.39] | 0.78(0.24)<br>[0.33-1.15] | t(40.31)= -0.90, *p* = .37,<br>95% CI [-0.25, 0.09] |
| Aperiodic exponent | 2.41 (0.20)<br>[1.95-2.84] | 2.42 (0.23)<br>[1.95-2.84] | 2.39 (0.17)<br>[1.97-2.67] | t(42.37)= -0.48, *p* = .64,<br>95% CI [-0.15, 0.09] |
| Communicative abilities (CSBS-DP) | 35.82(6.53)<br>[14-50] | 34.50 (7.47)<br>[14-48] | 37.04 (5.28)<br>[28-50] | t(40.81)= 1.37, *p* = .18,<br>95% CI [-1.21, 6.29] |

*Note.* Table 2 presents the descriptive statistics for neonatal aperiodic offset and exponent, and communicative skills at 12 months (CSBS-DP Infant-Toddler Checklist) for the total sample (*n* = 50) and separately by sex as *M* (SD) [range]. All variables were normally distributed, and t-tests indicated no significant sex differences.

### Aperiodic offset and communicative abilities

The multiple linear regression model, which included the linear and the quadratic effect of the neonatal aperiodic offset as well as infant sex, reached statistical significance (*R*^2^_adj_ = .12, *p* = .03), with only the quadratic term of the neonatal offset (β = −0.24, 95% CI [−0.47, −0.02], *p* = .04), but not the linear term (β = 0.16, 95% CI [−0.12, 0.44], *p* = .25) nor infant sex(β = −0.26, 95% CI [−0.83, 0.31], *p* = .36) significantly predicting communicative skills at 12 months. Although this model provides first evidence for linking aperiodic offset to communicative abilities, this finding was not replicated in the sensitivity analysis using robust regression (see SI, Table S3a), suggesting that the observed association may be sensitive to influential cases and warrants cautious interpretation.

### Aperiodic exponent and communicative abilities

The multiple linear regression model including the neonatal aperiodic exponent as both linear and quadratic terms, along with infant sex, significantly predicted subsequent communicative abilities (*R*^2^_adj_ = .14, *p* = .02). In this model, only the quadratic term of the aperiodic exponent (β = −0.27, 95% CI [−0.48, −0.06], *p* = .01) was significantly associated with infant communicative skills at 12 months, and neither infant sex (β = −0.26, 95% CI [−0.80, 0.29], *p* = .35) nor the linear term of the aperiodic exponent (β = 0.14, 95% CI [−0.13, 0.41], *p* = .31) showed a significant association, suggesting a non-linear relationship between neonatal aperiodic exponent and infant communicative skills at 12 months, confirmed by robust regression analysis (see SI, Table S3b). Specifically, the pattern was consistent with an inverted U-shaped association, with both lower and higher aperiodic exponent values related to lower communicative skills at 12 months (see Figure 4).

**Figure 4.**
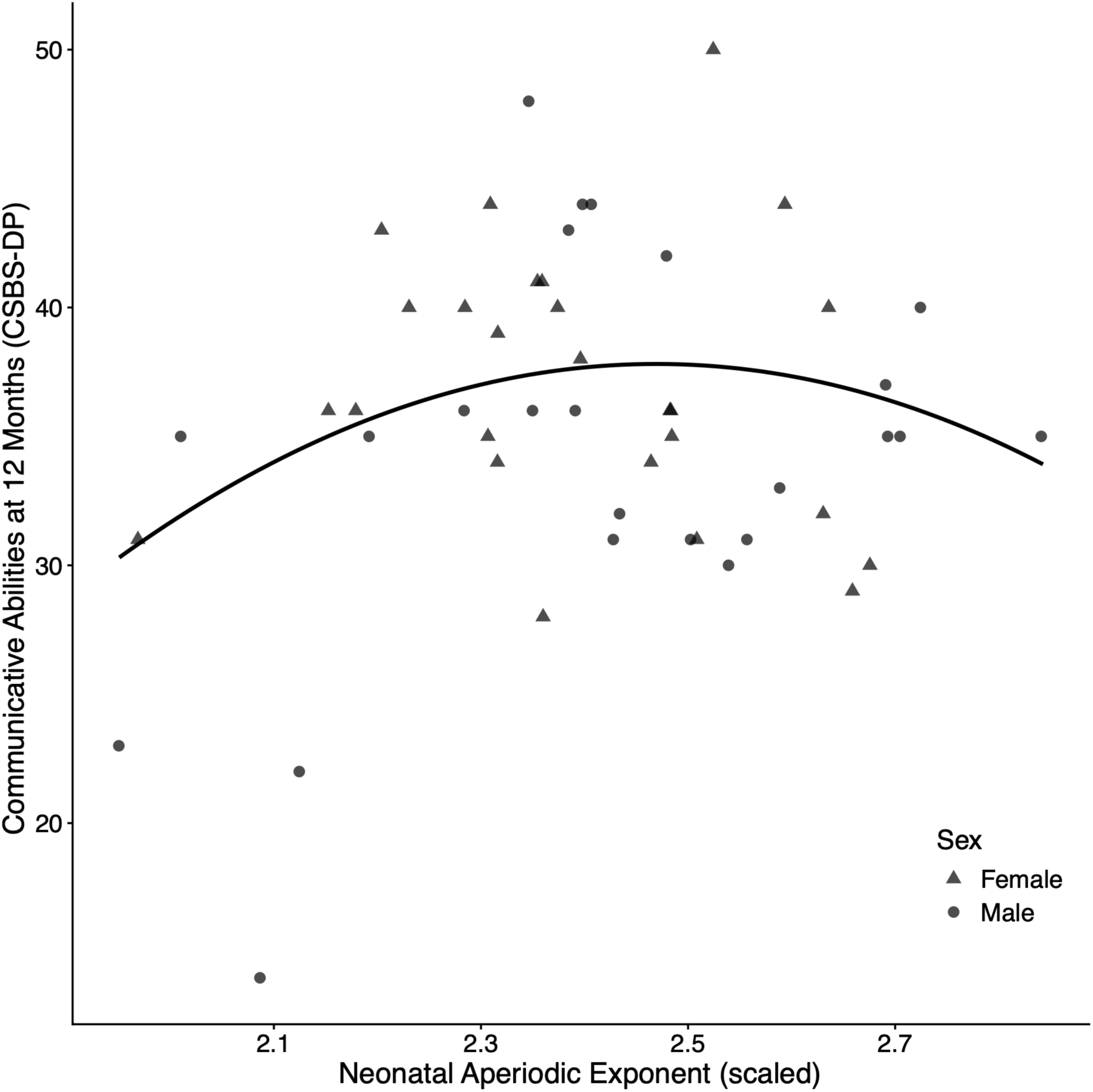
Inverted U-shaped association between neonatal aperiodic exponent and communicative abilities at 12 months (CSBS-DP Infant-Toddler Checklist), such that both lower and higher exponent values were associated with lower CSBS-DP scores (data jittered to prevent overlapping points, *n* = 50).

Control analyses limited to EEG periods, during which the infants were asleep, confirmed the predictive value of the neonatal aperiodic exponent for communicative abilities at 12 months in an inverted U-shaped pattern (see SI, Table S5b), while no effect was found for the aperiodic offset (see SI, Table S5a). Furthermore, gestational age at birth did not emerge as a significant predictor of later communication skills (see SI, Table S4), and thus does not account for the observed findings. Similarly, neither the absolute delta nor the theta power model reached statistical significance (see SI, Tables S6a and S6b).

## Discussion

In the present study, we investigated whether aperiodic activity at two weeks of age during EEG resting-state recording predicted later communicative abilities at 12 months. Our findings highlight neonatal aperiodic activity as a neurophysiological marker for communicative development. Specifically, the neonatal aperiodic exponent, potentially reflecting E/I balance, was found to be related to communicative skills at 12 months in an inverted U-shaped manner. That is, lower and higher values of the aperiodic exponent were found to relate to lower language abilities, and mid-range values to higher language abilities. In contrast, no robust association was found between the aperiodic offset and later communicative abilities in our data. Overall, our results suggest that aperiodic activity may play a role in early communicative development beyond previous clinical reports of associations with neurodevelopmental disorders.

Importantly, our data show a non-linear, inverted U-shaped relationship between the aperiodic exponent and subsequent communication skills, indicating that favorable developmental outcomes may be achieved at intermediate levels of the aperiodic exponent, rather than with linearly increasing or decreasing values. While empirical evidence linking aperiodic activity to language-related outcomes in early childhood remains limited, our findings can be viewed within the dynamic network communication framework [49], which discusses neural communication in the context of cognition, development, aging, and psychiatric and neurological disorders. Within this framework, deviations towards excessively steep or excessively flat aperiodic 1/f slopes may both reflect dysfunctional neural communication. Similarly, a systematic review highlighted substantial heterogeneity in reports of aperiodic activity across 38 neurological and psychiatric disorders, with findings almost evenly split between studies reporting an increased aperiodic exponent, decreased aperiodic exponent, or no difference in the clinical compared to the non-clinical group [61]. Evidence that altered exponents in either direction may be associated with atypical brain functioning further hints at a non-linear relationship between aperiodic activity and developmental outcome. Our findings reveal that such a non-linear pattern may also be present in a non-clinical infant sample. Here, intermediate levels of the aperiodic exponent may reflect a more favorable neurodevelopmental state, whereas deviations in either direction may indicate less optimal conditions for later development. Further research in non-clinical samples, particularly in early development, will need to establish whether the observed non-linear association reflects a reliable developmental pattern.

Critically, our findings further support the view that aperiodic neural activity should not be dismissed as background noise, but rather captures physiologically meaningful aspects of brain function. So far, the aperiodic offset has received considerably less attention compared to the exponent (see also [61]). In the present study, exponent and offset were not equally informative with respect to later communicative development; only the aperiodic exponent predicted subsequent communication abilities in an inverted U-shaped pattern, but this association was not confirmed in robustness analyses for the aperiodic offset. This pattern may suggest that, although offset and exponent are closely intertwined (see also [22]), the exponent represents a more sensitive marker of subsequent communicative abilities. Mechanistically, the aperiodic exponent has been linked to E/I balance [28], with current evidence largely based on computational models and intracranial electrophysiological recordings. E/I balance, in turn, is thought to support robust and efficient information processing [62,63]. Balanced excitation and inhibition enable neural populations to respond reliably to relevant sensory input while maintaining selectivity and preventing excessive spread of activity [62,63] (for a review, see [30]). When excitation dominates, cortical activity may become less precisely regulated, potentially reducing neural response-selectivity and compromising stimulus encoding [62,64]. This may be particularly relevant to social and cognitive functioning, as animal evidence indicates that experimentally increased excitation specifically in prefrontal cortical excitatory neurons can impair these domains [65]. Conversely, excessive inhibition may decrease cortical responses to incoming input, thereby limiting responsiveness and constraining information processing [64] (for a review, see [30]). Importantly, E/I balance does not appear static throughout development, but changes as neural circuits mature (for an overview, see [66]). Given the role of E/I balance in regulating neural responsiveness and information processing, developmental variation in the aperiodic exponent may therefore provide a window into neural dynamics relevant to cognitive and communicative functioning.

In the context of early neurodevelopment, the aperiodic exponent may offer insight into the functional maturation of developing neural circuits [33]. Developmental animal research has shown that increasing inhibition drives the decorrelation of neural activity, which is thought to reflect more efficient information processing [33]. Similar patterns in neonatal EEG [33] suggest that developmental E/I imbalances may be relevant to neurodevelopmental outcomes [33,67]. By focusing on variation in early communicative abilities within a non-clinical sample, our study extends previous findings on aperiodic activity and developmental outcomes, which were largely confined to neurodevelopmental disorders, genetic syndromes, or an elevated risk of developmental disorders (for a review, see [68]). While this evidence does not allow for direct comparison, it nonetheless offers tentative support for links between aperiodic activity and language-related outcomes. Specifically, developmental trajectories in aperiodic neural activity have been linked to language abilities in infants with and without familial risk for autism [46] and in children aged six to 17 years with and without ASD [47], while no relationship was found for children with Down syndrome [42]. Critically, our findings highlight that aperiodic neural activity may be informative not only in children with developmental difficulties but also for understanding individual differences in communication in typically developing children, although further research is needed to understand how these neurobiological processes translate into later observable communicative behavior.

By showing that aperiodic activity can offer further insight into early developmental variation, our findings broaden recent approaches to predicting language development from neural activity, which have so far mainly investigated oscillatory activity using traditional frequency-band power measures (e.g., [17,18]). Within our neonatal sample, additional analyses of conventional delta and theta power did not reveal associations with later communicative abilities, suggesting that aperiodic activity may capture aspects of neural variation not readily reflected in conventional band-power measures. Taken together, these considerations highlight the potential value of separating aperiodic and periodic components for understanding early neural mechanisms (and how they relate to observable behavior), particularly as the spectral composition of neural activity changes markedly across infancy [32,35,36]. This approach enables more precise attributions of language-related abilities to different neural signals and their interplay, thereby offering a more differentiated perspective on the neural dynamics supporting early communication [22]).

Although our study benefits from a prospective longitudinal design and a comparatively large sample size, several methodological limitations should be acknowledged. Due to the very early assessment, with EEG data collected as early as two weeks after birth, most recordings were conducted in the families’ homes. Despite less standardization than in lab settings, EEG recordings followed a fixed procedure, and this flexible approach encouraged participation by reducing the burden on families and providing a more natural setting during this sensitive period. In addition, the relatively homogeneous demographic composition of the sample may limit the generalizability of the present findings. Further research should therefore include more socioeconomically (e.g., [69]) and demographically diverse samples. As is common in neonate EEG research, data collection can be complicated by fluctuations in infant behavioral state, crying, fussiness, and variation in sleep-wake state (for a methodological overview, see [70]). Nevertheless, the relatively long EEG resting-state recordings (over 12 minutes), together with a multistage exclusion procedure, ensured sufficient artifact-free data required for the data analysis. To address variability related to sleep-wake state and evaluate the robustness of the findings, additional analyses were restricted to infants sleeping during data collection, confirming the non-linear association between the aperiodic exponent and subsequent communicative abilities. Moreover, instead of examining language abilities directly, we focused on communicative abilities, as these capture precursor skills that underlie language development [48]. The CSBS-DP Infant-Toddler Checklist is a well-established screening measure of early communication abilities, not intended for clinical diagnosis; therefore, scores should be interpreted as reflecting variation in early communicative behavior rather than clinical classification.

Communication is an essential skill in early social interaction, building the foundation for emerging language development. Our study provides the first evidence that aperiodic neural activity in neonates relates to their later communicative skills. Our findings suggest that an intermediate level of the aperiodic exponent at birth may be optimal for early communicative development, consistent with a balanced E/I state. Taken together, these findings broaden the perspective on aperiodic components of neural activity by suggesting that the aperiodic exponent, in particular, may be informative not only as a potential biomarker in neurodevelopmental disorders but also for capturing variability in early development.

## Ethical Information

Before participation, mothers provided informed consent, and the ethics committee of the Paris Lodron University of Salzburg approved the experimental procedure (EK-GZ: 12/2013, positive vote on 23^rd^ of January, 2014, Addendum: 17^th^ November, 2021).

## Acknowledgements

The authors would like to thank the mothers, fathers, and children who took part in this project, as well as the students who helped collect the data.

## Data availability

Non-sensitive data is available from the corresponding author upon reasonable request.

