## Supplementary Information for "Talking Brains: Neonatal Aperiodic Activity in Resting State EEG Relates to Later Communicative Abilities"

### SUPPLEMENTARY MATERIAL

### SUPPLEMENTARY METHODS

#### S1 EEG layout and region of interest

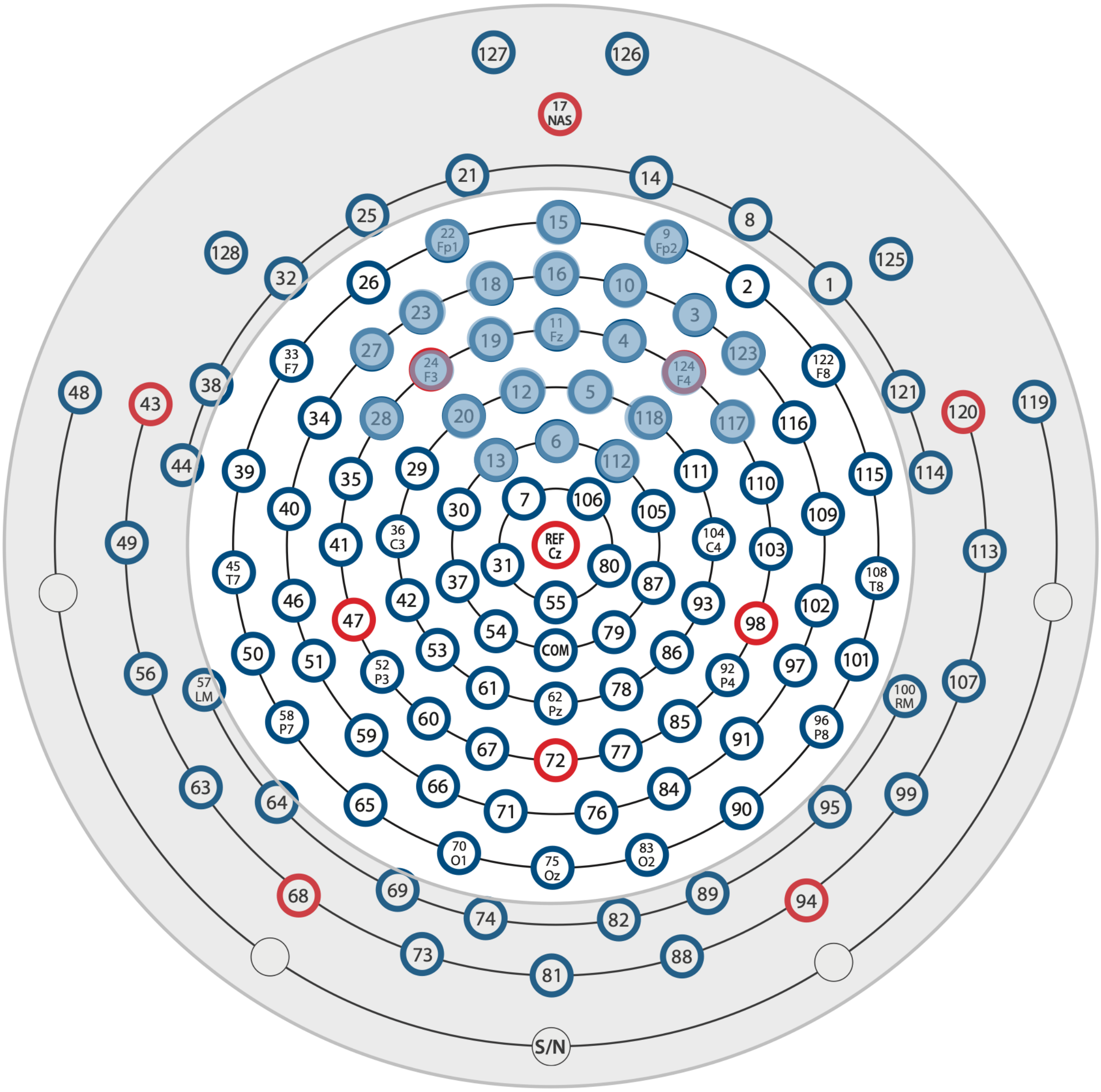


Figure S1. Illustration of the EEG electrode net used. Preprocessing was performed using all inner electrodes, excluding the two outermost rings, highlighted in grey. For statistical analysis, we focused on *n* = 24 electrodes in the frontal region, highlighted in blue. Figure adapted from the Healthy Brain/FCON 1000 website, available online at http://fcon_1000.projects.nitrc.org/indi/cmi_healthy_brain_network/File/_eeg/EEG_128_channel_array_map.pdf and based on the dataset described by Alexander et al. [1].

### S2 Comparison of FOOOF fits across 1-15 Hz and 1-30 Hz

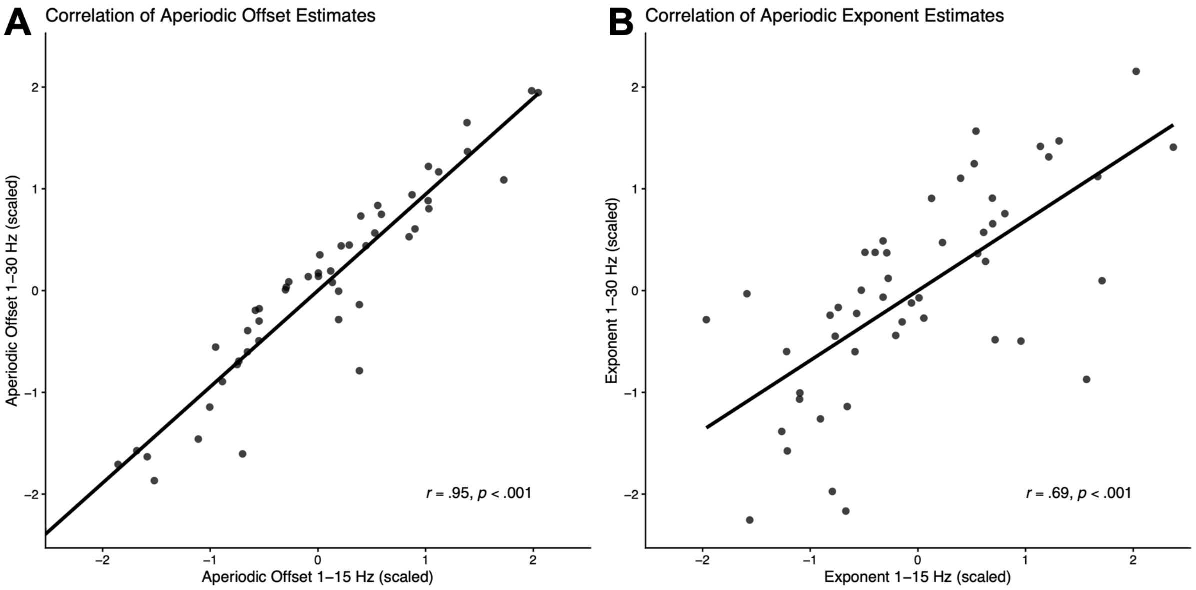


Figure S2. Pearson correlations between (A) aperiodic offset ($r$ = 0.95, $p < .001$) and (B) exponent ($r$ = 0.69, $p < .001$) estimated using FOOOF across 1–15 Hz and 1–30 Hz. Based on this strong agreement, we selected the 1–30 Hz range for all subsequent analyses, following recommendations that broader fitting ranges improve the robustness and interpretability of aperiodic parameter estimates (see [2]).

### SUPPLEMENTARY ANALYSES

#### S3 SENSITIVITY ANALYSES USING ROBUST REGRESSION

**Table S3a: Sensitivity analysis using robust regression for the association between neonatal aperiodic offset and communicative abilities at 12 months (*n* = 50)**

| | β | 95% CI | *p* |
|---|---|---|---|
| Aperiodic offset (linear) | 0.13 | [-0.45, 0.71] | .66 |
| Aperiodic offset (quadratic) | -0.17 | [-0.75, 0.41] | .55 |
| Sex | -0.25 | [-0.86, 0.35] | .40 |

*Note.* Using robust regression analysis, the aperiodic offset does not show any predictive power for explaining communicative abilities at 12 months ($R^2_{adj}$ = 0.04).

**Table S3b: Sensitivity analysis using robust regression for the association between neonatal aperiodic exponent and communicative abilities at 12 months (*n* = 50)**

| | β | 95% CI | *p* |
|---|---|---|---|
| Aperiodic exponent (linear) | 0.06 | [-0.28, 0.40] | .72 |
| Aperiodic exponent (quadratic) | -0.23 | [-0.42, -0.05] | .02 |
| Sex | -0.17 | [-0.77, 0.43] | .57 |

*Note.* Robust regression analysis confirmed the predictive value of the aperiodic exponent on later communicative abilities ($R^2_{adj}$ = 0.08).

#### S4 SENSITIVITY ANALYSIS FOR GESTATIONAL AGE

Because the aperiodic exponent was related to gestational age at birth, we performed a sensitivity analysis in which we replaced the aperiodic component with gestational age at birth. Our results indicate that gestational age did not predict later communicative skills at 12 months, and the overall model did not reach statistical significance.

**Table S4: Sensitivity analysis: Multiple regression analysis examining the association between gestational age at birth and communicative abilities at 12 months (*n* = 50)**

| | β | 95% CI | *p* |
|---|---|---|---|
| Gestational age (linear) | -0.10 | [-0.49, 0.28] | .59 |
| Gestational age (quadratic) | 0.04 | [-0.16, 0.23] | .71 |
| Sex | -0.45 | [-1.07, 0.17] | .15 |

*Note.* The regression model, which included linear and quadratic terms for gestational age and infant sex, was not statistically significant in predicting communicative abilities at 12 months ($R^2_{adj}$ = 0.00, $p$ = .38).

### S5 SENSITIVITY ANALYSIS IN SLEEPING NEONATES

#### Sleep-wake state coding

To differentiate sleep and wake periods, a trained infant sleep-staging expert manually evaluated the EEG files in 30-second segments according to the American Academy of Sleep Medicine's (AASM) guidelines for children (version 2.1, 2.2) [3,4]. Sleep-wake classification was based on EEG, respiration activity measured using a respiratory belt, and movement patterns observed in the video recordings. Of the 50 participants, 45 participants had at least 100 clean trials obtained during sleep ($M_{\text{clean epochs}}$ = 352.11, $SD$ = 137.40, range = 106-612), and were thus included in the analysis (aperiodic offset: $M$ = 0.85, $SD$ = 0.29, range = [0.28-1.48]; aperiodic exponent: $M$ = 2.44, $SD$ = 0.21, range = [1.95, 2.85]).

**Table S5a: Sensitivity Analysis: Multiple regression analysis examining the association between the neonatal aperiodic offset and communicative abilities at 12 months in sleeping infants (*n* = 45)**

| | β | 95% CI | *p* |
|---|---|---|---|
| Aperiodic offset (linear) | 0.11 | [-0.21, 0.42] | .50 |
| Aperiodic offset (quadratic) | -0.14 | [-0.38, 0.11] | .28 |
| Sex | -0.24 | [-0.87, 0.39] | .44 |

*Note.* In the sleeping subsample, no significant effect of neonatal aperiodic offset on communicative abilities at 12 months was observed ($R^2_{adj}$ = 0.00, $p$ = .44).

**Table S5b: Sensitivity Analysis: Multiple regression analysis examining the association between the neonatal aperiodic exponent and communicative abilities at 12 months in sleeping infants ($n$ = 45)**

| | β | 95% CI | $p$ |
|---|---|---|---|
| Aperiodic exponent (linear) | 0.05 | [-0.25, 0.35] | .72 |
| Aperiodic exponent (quadratic) | -0.27 | [-0.50, -0.05] | .02 |
| Sex | -0.17 | [-0.75, 0.42] | .57 |

*Note.* In the sleeping subsample, the significant quadratic effect of the neonatal aperiodic exponent confirmed the inverted U-shaped association found for the full sample with communicative abilities at 12 months, although the overall model narrowly missed conventional significance ($R^2_{adj}$ = 0.10, $p$ = .06).

### S6 SENSITIVITY ANALYSIS USING ABSOLUTE DELTA AND THETA POWER

#### Conventional spectral power

To examine whether the observed associations were also reflected in conventional spectral power, absolute delta (1- <3 Hz) and theta (3- 6 Hz) power were calculated for the same participants using the same retained, preprocessed EEG data as in the primary analyses. Power spectra were estimated for each electrode using a Hanning-tapered Fast Fourier Transform over the 1-30 Hz range at 0.25-Hz intervals and averaged across epochs. Power was integrated within each frequency band using the trapezoidal rule and subsequently averaged across $n$ = 24 electrodes within the frontal region of interest (delta: $M$ = 4.18, $SD$ = 2.49, range = [0.87-11.81]; theta: $M$ = 1.23, $SD$ = 0.75, range = [0.29-4.63]). Because distributions were right-skewed (Shapiro-Wilk test; delta: $W$ = 0.85, $p$ < .001; theta: $W$ = 0.84, $p$ < .001), delta and theta power were log10-transformed and scaled prior to statistical analyses. Hierarchical regression models were fitted separately for delta and theta power, including infant sex in all models. First, we examined linear associations, consistent with previous spectral power findings in infants (e.g., [5]). In a second step, quadratic terms were added to allow direct comparison with the nonlinear association observed for the aperiodic exponent. Please note that the quadratic model examining delta power showed increased VIF values (<10), indicating potential multicollinearity.

**Table S6a: Sensitivity Analysis: Multiple regression analysis examining the association between neonatal absolute delta power and communicative abilities at 12 months supplemented by robust regression analyses (*n* = 50)**

| | | | | Robust Regression [RR] | | |
|---|---|---|---|---|---|---|
| | β | 95% CI | *p* | β | 95% CI | *p* |
| **Linear model** | | | | | | |
| Delta power (linear) | 0.15 | [-0.13, 0.44] | 0.28 | -0.06 | [-0.44, 0.31] | .75 |
| Sex | -0.43 | [-0.99, 0.14] | 0.14 | -0.20 | [-0.81, 0.40] | .50 |
| **Quadratic model** | | | | | | |
| Delta power (linear) | 0.13 | [-0.15, 0.42] | .06 | -0.05 | [-0.85, 0.74] | .89 |
| Delta power (quadratic) | -0.24 | [-0.52, 0.04] | .10 | -0.01 | [-0.86, 0.84] | .98 |
| Sex | -0.36 | [-0.92, 0.20] | .20 | -0.21 | [-1.04, 0.62] | .61 |

*Note.* No significant effect of neonatal absolute delta power on communicative abilities at 12 months was observed (Linear model: $R^2_{adj}$ = 0.02 [RR: -0.02], $p$ = .22; Quadratic model: $R^2_{adj}$ = 0.06 [RR:-0.04], $p$ = .12).

**Table S6b: Sensitivity Analysis: Multiple regression analysis examining the association between neonatal absolute theta power and communicative abilities at 12 months supplemented by robust regression analyses (*n* = 50)**

| | | | | Robust Regression [RR] | | |
|---|---|---|---|---|---|---|
| | β | 95% CI | *p* | β | 95% CI | *p* |
| **Linear model** | | | | | | |
| Theta power (linear) | 0.21 | [-0.07, 0.49] | 0.14 | 0.03 | [-0.42, 0.48] | .90 |
| Sex | -0.39 | [-0.95, 0.17] | 0.16 | -0.25 | [-0.79, 0.30] | .37 |
| **Quadratic model** | | | | | | |
| Theta power (linear) | 0.20 | [-0.09, 0.48] | .12 | 0.04 | [-0.64, 0.71] | .91 |
| Theta power (quadratic) | -0.17 | [-0.46, 0.11] | .22 | 0.00 | [-0.63, 0.64] | .99 |
| Sex | -0.35 | [-0.91, 0.21] | .21 | -0.25 | [-0.89, 0.38] | .43 |

*Note.* No significant effect of neonatal absolute theta power on communicative abilities at 12 months was observed (Linear model: $R^2_{adj}$ = 0.04 [RR: -0.02], $p$ = .13; Quadratic model: $R^2_{adj}$ = 0.05 [RR: -0.04], $p$ = .14).
